# Protein large language model assisted one-to-one gene homology mapping in cross-species single-cell transcriptome integration

**DOI:** 10.1101/2025.10.17.683009

**Authors:** Ze-Yu Kuang, Yuan-Chen Sun, Na-Na Wei, Yu-Juan Wang, Hua-Jun Wu

## Abstract

Cross-species integration of single-cell transcriptomes requires establishing gene correspondences to enable comparative analysis of expression profiles across organisms. Current approaches predominantly rely on Ensembl homology tables; although gene-family expansion and contraction can reflect biologically meaningful evolutionary divergence, default many-to-many mappings can overweight expanded gene-family signals during integration and generate mapping-associated micro-clusters that lack clear cell-type identity, thereby complicating direct cell-type alignment. While restricting mappings to a one-to-one scheme suppresses such artifacts, it reduces the number of homology gene pairs by approximately 8% (∼900 pairs). To address this limitation, we develop a protein large language model (pLLM)-based gene homology mapping strategy that boosts the number of homology gene pairs. By integrating pLLM-derived representations with sequence similarity, we construct a fused mapping approach, which achieves top performance in a comprehensive benchmark based on a curated cross-species atlas—spanning nine datasets, 11 species, and over 3.2 million cells. Our method further identifies previously unannotated cell-type marker pairs, facilitating novel cross-species marker discovery. These results establish a robust framework for gene homology mapping in cross-species transcriptome integration, improving both accuracy and biological interpretability.

## Introduction

The rapid proliferation of single-cell transcriptome atlases across diverse species has deepened our understanding of species-specific and conserved cell types and their molecular programs (Tosches et al. 2018; Han et al. 2022; Zhang et al. 2025). Joint analysis of homologous tissues from different organisms enables the dissection of lineage and functional conservation at evolutionary scales and supports comparative studies at higher taxonomic levels (Jorstad et al. 2023; Rosen et al. 2024).

Cross-species integration hinges on two steps: (1) gene homology mapping (Emms and Kelly 2019; Song et al. 2023)—mapping genes across species to a shared index so that expression matrices can be compared—and (2) correction of batch/species effects within the mapped homology genes (Luecken et al. 2022; Zhu et al. 2023). While substantial progress has been made on the second step, the first—how to map genes across species—remains underexplored. Many studies rely on Ensembl homology tables (Breschi et al. 2017; Shafer 2019; Liu et al. 2023); however, default many-to-many mappings can conflate gene-family expansion with mapping uncertainty. In the context of direct cross-species cell-type alignment, such mappings may overweight expanded gene-family signals and thereby complicate the biological interpretation of downstream integration.

Protein large language models (pLLMs) provide an opportunity to rethink cross-species gene homology mapping (Lin et al. 2023; Abramson et al. 2024; Hayes et al. 2025). Trained solely on primary amino-acid sequences, these models learn high-dimensional embeddings that are comparable across species, potentially bypassing name inconsistencies and directly capturing functional similarity. Building on this idea, we complement classical homology with ESM2-derived gene embeddings and develop a set of one-to-one–oriented mapping strategies based on: (1) Ensembl sequence identity and orthology confidence (HM_O2O), (2) ESM2 cross-species embedding correlations (LM_O2O), and (3) a fused one-to-one scheme that integrates both sources of evidence (HL_O2O).

Here, we aim to systematically evaluate whether protein-language-model-derived gene representations can improve cross-species gene homology mapping for single-cell transcriptome integration. Using a multi-species benchmark spanning diverse tissues and evolutionary distances, we compare many-to-many, sequence-based one-to-one, embedding-based one-to-one, and fused one-to-one mapping strategies, with the goal of assessing how different gene-mapping choices affect integration performance and the identification of putative cross-species marker-gene correspondences.

## Results

### pLLM-based one-to-one gene homology mapping improves integration of single-cell transcriptomics

A core challenge in cross-species single-cell transcriptome integration is to mapping homology genes across species (Tarashansky et al. 2021), which requires establishing a gene homology mapping to concatenate and align expression matrix under a same homology-gene-space (Rosen et al. 2024).

Previous methods have primarily utilized Ensembl homology tables (Herrero et al. 2016), employing either the default many-to-many mappings (ENS_M2M) or a filtered set of “ortholog_one2one” entries (ENS_O2O). These approaches, however, are fundamentally constrained by sequence similarity and may miss ortholog pairs that have large phylogenetic distance (Kilinc et al. 2023). To overcome this limitation, we introduced a strategy that leverages the protein language model ESM2 to learn cross-species gene representations. This enables the data-driven discovery of functionally analogous genes independent of existing annotation or nomenclature constraints. We refer to this mapping as LM_O2O, that establishes global one-to-one mappings through a greedy selection procedure that prioritizes correlations between cross-species gene embeddings (Fig. 1A, see Methods).

**Fig. 1.**
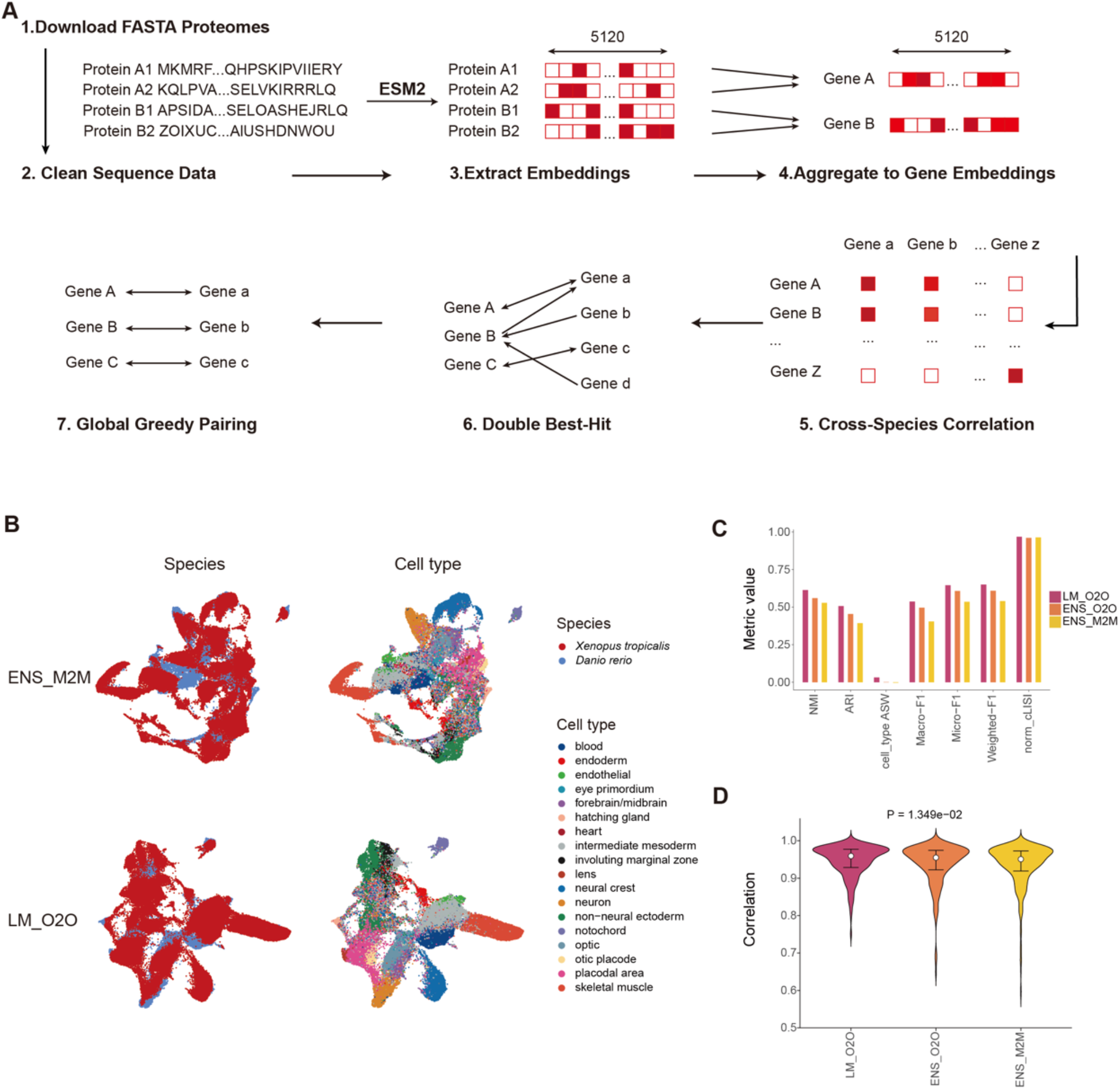
pLLM-based cross-species gene homology mapping improves integration of single-cell transcriptomics. **A**, Workflow of pLLM-based one-to-one (LM_O2O) gene mapping. Protein sequences from two species are used to produce one-to-one gene correspondences for downstream cross-species single-cell transcriptome integration. **B**, UMAP visualization of Embryo dataset integration using Ensembl many-to-many mappings (ENS_M2M, left) versus LM_O2O (right), colored by species (top) and cell type (bottom). **C**, Performance comparison of ENS_M2M, ENS_O2O (Ensembl ortholog_one2one), and LM_O2O across seven integration metrics: NMI, ARI, Cell_type_ASW, Macro_F1, Micro_F1, Weighted_F1, and normalized cLISI. **D**, Distribution of correlation coefficients for gene pairs identified by each method. Statistical significance (P = 1.349×10^−2; two-sided test) is indicated for the comparison between ENS_M2M and LM_O2O.

On an Embryo (Emb) (Wagner et al. 2018) dataset, LM_O2O achieves superior species mixing and clearer cell-type separation compared to ENS_M2M (Fig. 1B), with particularly improvements in skeletal, neural crest, blood, and intermediate mesoderm cell populations. A benchmark across seven clustering metrics (NMI, ARI, Cell_type_ASW, Macro-F1, Micro-F1, Weighted-F1, and norm_cLISI) confirms that LM_O2O outperformed both ENS_O2O and ENS_M2M (Fig. 1C). Furthermore, in contrast to LM_O2O, both ENS_M2M and ENS_O2O exhibit a higher proportion of gene pairs with low ESM2-based correlation (Fig. 1D), indicating the inclusion of more putative unreliable alignments that likely introduce noise. Together, these results indicate that ESM2-driven one-to-one mapping uncovers novel and effective cross-species correspondences, leading to more accurate integration of cross-species single-cell transcriptomes.

### Benchmarking different gene homology mapping strategies using a cross-species transcriptome atlas

To assess the generalizability of LM_O2O gene homology mapping approach across diverse tissues and species, we assembled nine datasets spanning 11 species comprising over 3.2 million cells (Fig. 2A–B). In addition to the two Ensembl baselines (ENS_M2M and ENS_O2O), we developed a homology-based one-to-one strategy (HM_O2O; see Methods). This method establishes global one-to-one mappings through a greedy selection procedure that prioritizes Ensembl orthology confidence and sequence identity. Benchmarking revealed that HM_O2O and LM_O2O achieve broadly comparable integration quality while capturing partially distinct sets of gene pairs (Supplemental Fig. S1A, B). To leverage this complementarity, we merged their candidate pairs and re-enforced a global one-to-one constraint, resulting in a fused mapping strategy termed HL_O2O (see Methods). This approach integrates both sequence similarity and protein language model representations, and it includes the highest number of unique one-to-one mapping gene pairs among the O2O strategies (Supplemental Fig. S2). Compared with HM_O2O, HL_O2O introduced an average of 512 additional gene pairs per dataset, enriching the homologous information available for cross-species gene matching while preserving the one-to-one constraint (Supplemental Fig. S1C).

**Fig. 2.**
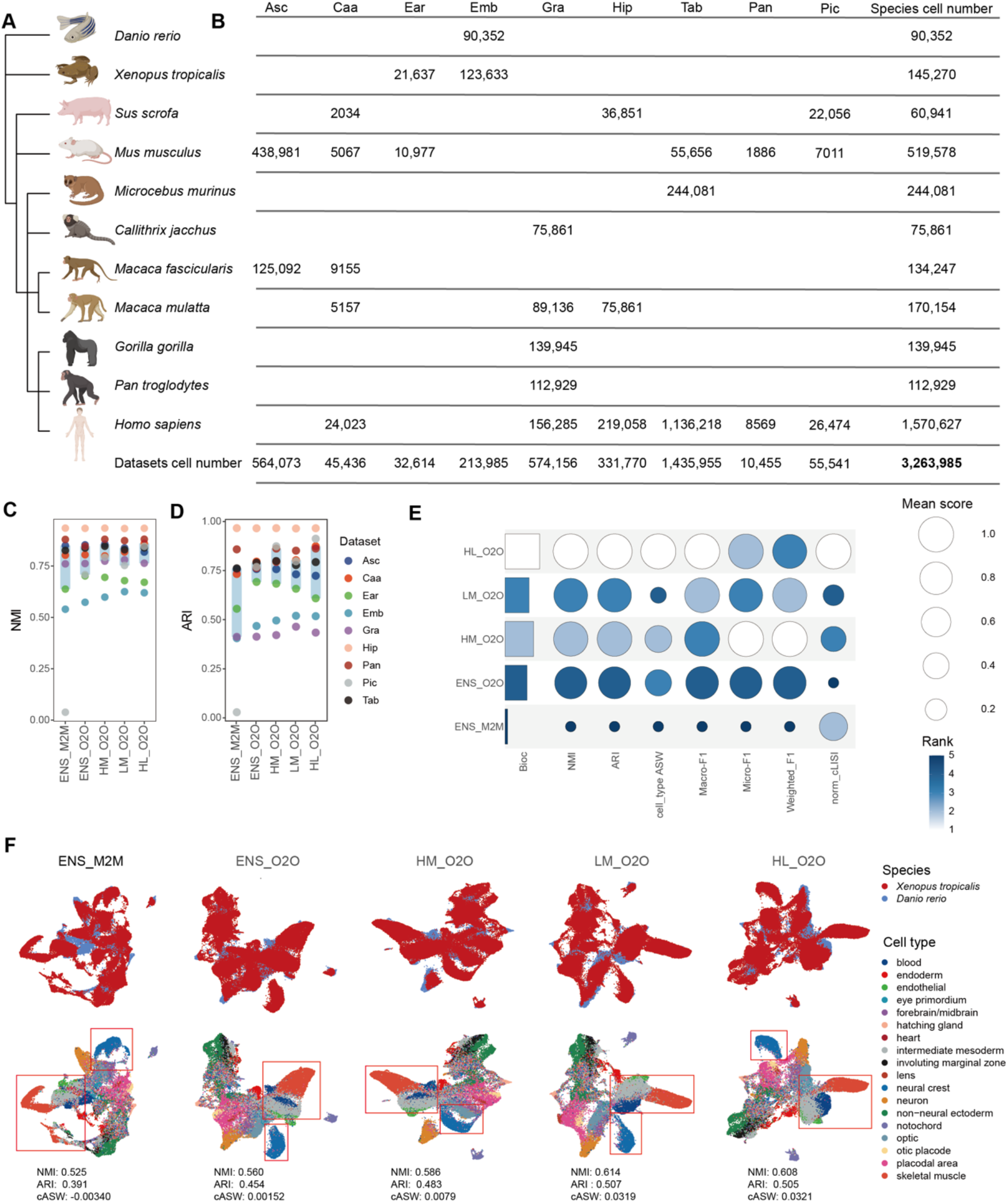
Benchmarking cross-species gene homology mapping across diverse datasets. **A**, Phylogenetic overview of species included in the benchmark. **B**, Summary of the number of cells across species and tissues. Asc (A single cell map of antisense oligonucleotide activity in the brain), Caa (Cell Atlas of Aqueous Humor), Ear (The single cell atlas of inner ear)A, Emb (Embryo), Gra (Post-mortem cortical tissue of Great apes), Hip (Hippocampus), Pan (Pancreas), Pic (Pancreatic islet cells), Tab (Tabula Sapiens, Tabula Microcebus and single cells from different organs of mice). **C–D**, Distribution of normalized mutual information (NMI; c) and adjusted rand index (ARI; d) values across the nine datasets for all five mapping strategies. Light-blue boxes represent the interquartile range (IQR). **E**, Bubble heatmap showing, for each method, the mean performance per dataset across seven clustering metrics (NMI, ARI, Cell_type_ASW, Macro_F1, Micro_F1, Weighted_F1, normalized cLISI) and the aggregated biological conservation score (Bioc). Bubble size and color intensity correspond to metric values and ranking, respectively. **F**, UMAP visualizations of Embryo dataset integration under the five mapping strategies, colored by species (top) and cell type (bottom); Key metrics (NMI, ARI and Cell_type_ASW) are annotated below each panel. Red rectangles highlight regions where where methodological differences in integration quality are most pronounced.

We then systematically evaluated the five strategies across all nine datasets using seven biology-conservation metrics: NMI, ARI, cell-type ASW, Macro-F1, Micro-F1, Weighted-F1, and normalized cLISI (Supplemental Table S1). Across the benchmark datasets, the most consistent performance gain was observed when moving from the many-to-many ENS_M2M strategy to one-to-one mapping strategies, including HL_O2O, LM_O2O, and HM_O2O (Fig. 2C–D; Supplemental Table S1). To facilitate an overall comparison, we normalized the per-dataset metric means and combined them into a unified biology conservation score (Bioc) (Song et al. 2023). HL_O2O achieved the highest Bioc (Fig. 2E), although HM_O2O and LM_O2O showed closely related performance across several core metrics (Supplemental Table S1), indicating that the fused strategy provides the best overall balance across metrics while retaining the main advantage of the one-to-one constraint. The improvement associated with the newly introduced one-to-one mapping approaches was also visually apparent in the Embryo dataset, where the HL_O2O UMAP projection showed strong species mixing and cell-type coherence (Fig. 2F). Notably, cell populations such as skeletal muscle, blood, and intermediate mesoderm show a clear separation pattern that aligns with the metric-based rankings, further validating the quantitative results.

### Many-to-many homology can overweight gene-family signals and produce mapping-associated micro-clusters

During cross-species integration, we observed dataset-specific mapping-associated micro-clusters when using the Ensembl many-to-many baseline (ENS_M2M). In both the pancreatic islet (Pic) (Tritschler et al. 2022) and cochlear (Ear) (Wang et al. 2024) datasets, UMAP visualizations colored by species and cell types revealed small, isolated clusters that were largely disconnected from major annotated cell-type manifolds when using ENS_M2M. These micro-clusters were substantially reduced by the one-to-one fusion strategy HL_O2O (Fig. 3A–B). Deconvolution of the cluster compositions showed that these micro-clusters had mixed cell-type identities rather than a single coherent annotation (Fig. 3C–D).

**Fig. 3.**
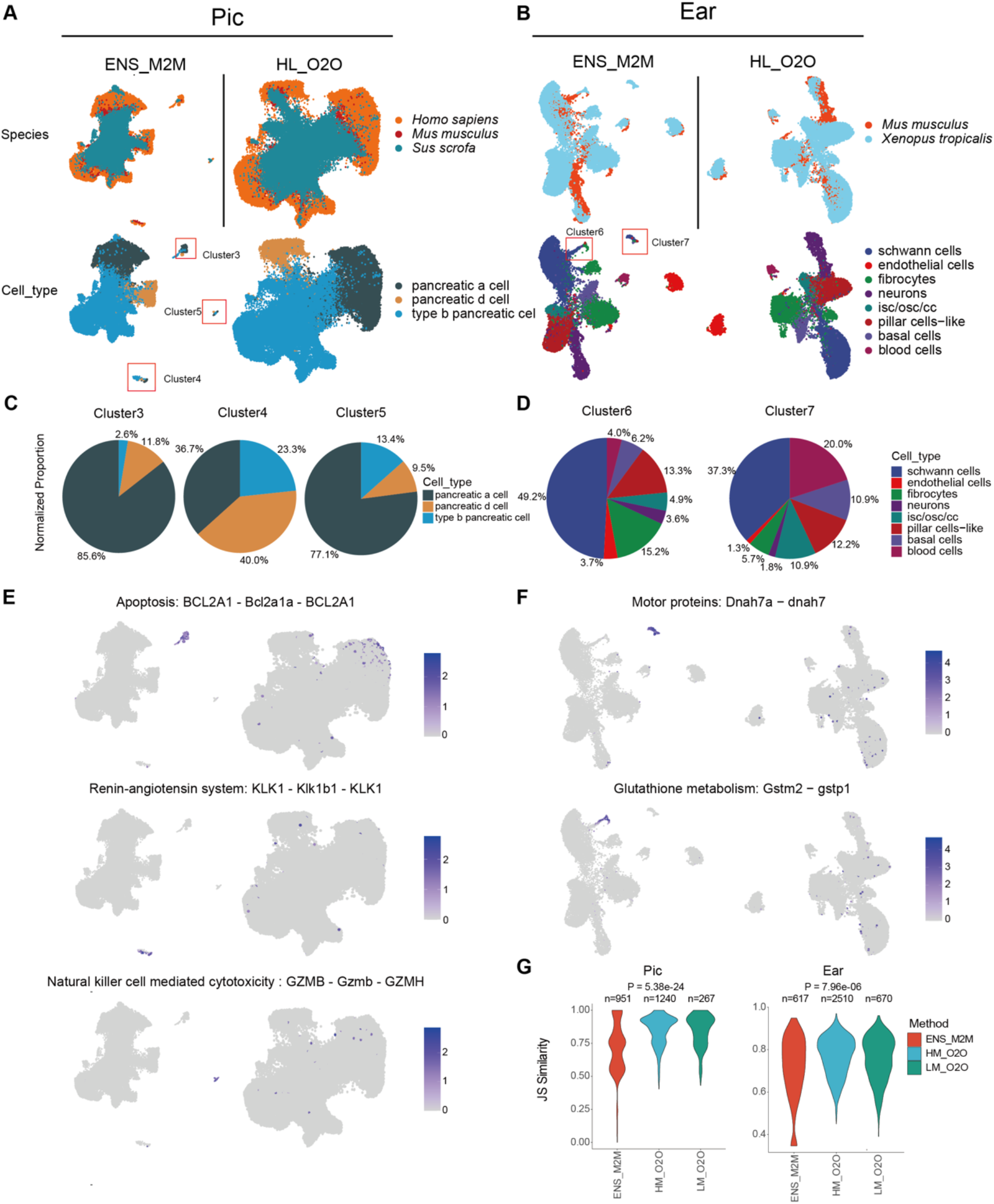
Many-to-many homology can overweight gene-family signals and produce mapping-associated micro-clusters during cross-species integration. **A–B**, UMAP visualizations of pancreatic islet and cochlear datasets, shown by species (top) and by cell type (bottom). Red boxes indicate micro-clusters observed under ENS_M2M that show mixed cell-type composition and reduced alignment with annotated cell types; results from the HL_O2O strategy are provided as a reference. **C–D**, Normalized cell-type composition analysis of ENS_M2M-specific micro-clusters: three clusters (3–5) in Pic (**C**) and two clusters (6–7) in Ear (**D**), revealing mixed cell-type identities. **E–F**, Expression patterns of representative marker genes from the micro-clusters, visualized on UMAP embeddings generated by ENS_M2M (left) and HL_O2O (right) in Pic (**E**) and Ear (**F**) datasets. **G**, Jensen-Shannon (JS) similarity of expression profiles for strategy-specific gene mappings in Pic and Ear datasets. ENS_M2M-unique pairs show systematically lower JS similarity compared to HM_O2O and LM_O2O. The P-value corresponds to a one-sided Wilcoxon test assessing whether ENS_M2M produces lower similarity than LM_O2O.

Marker inspection further indicated that these signals were enriched by specific gene families, whose influence may be disproportionately weighted by many-to-many mappings during integration. Notable examples in the Pic dataset include apoptosis (BCL2A1) (Vogler 2012), renin–angiotensin (KLK1) (Schmaier 2003), and NK-cell cytotoxicity (GZMB) (Kim et al. 2011); in the Ear dataset, examples include motor proteins (Dnah7a) (Dong et al. 2014) and glutathione metabolism (Gstm2) (Kumar and Reddy 2001). Feature plots comparing ENS_M2M and HL_O2O showed that these markers aggregate spuriously only with many-to-many mapping (Fig. 3E–F). This pattern is consistent with a mapping-overweighting effect when a single gene in one species is linked to multiple counterparts in another, which can disproportionately amplify gene-family-associated signals and complicate direct cross-species cell-type alignment.

To quantify the expression concordance of mapped gene pairs, we compared the Jensen–Shannon (Kumar and Reddy 2001) (JS) similarity of mappings uniquely identified by ENS_M2M, HM_O2O, and LM_O2O in the Pic and Ear datasets. ENS_M2M-specific pairs exhibited lower JS similarity than those from HM_O2O and LM_O2O, with the difference relative to LM_O2O being statistically significant (one-sided Wilcoxon test; Fig. 3G). These findings indicate that, for direct cross-species cell-type alignment, unconstrained many-to-many homologies can introduce mapping-associated ambiguity, whereas one-to-one strategies help reduce this ambiguity while retaining functionally informative cross-species correspondences.

### Novel marker-gene pair discovery by pLLM-based gene homology mapping

To evaluate the added value of ESM-embedding-based mapping beyond sequence-based homology, we focused on gene pairs for which LM_O2O diverged from HM_O2O or Ensembl-based annotations. These cases are particularly informative because they allow us to ask whether protein language model embeddings can recover functionally corresponding genes that are missed or ambiguously represented by sequence-based homology mappings. To characterize the properties of LM_O2O–derived cross-species mappings, we computed the JS similarity of mapped homology gene expression for three mapping sources (ENS_M2M, HM_O2O, and LM_O2O) across nine datasets (Fig. 4A). ENS_M2M exhibited consistently and significantly lower JS similarity than both one-to-one strategies in every dataset, supporting the rationale for enforcing a strict one-to-one constraint for reliable cross-species mapping. Notably, LM_O2O and HM_O2O performed comparably in all but two datasets, indicating that language model-derived embeddings not only align with sequence homology-based evidence but may also complement it by recovering biologically meaningful gene pairs that may be overlooked by sequence similarity alone.

**Fig. 4.**
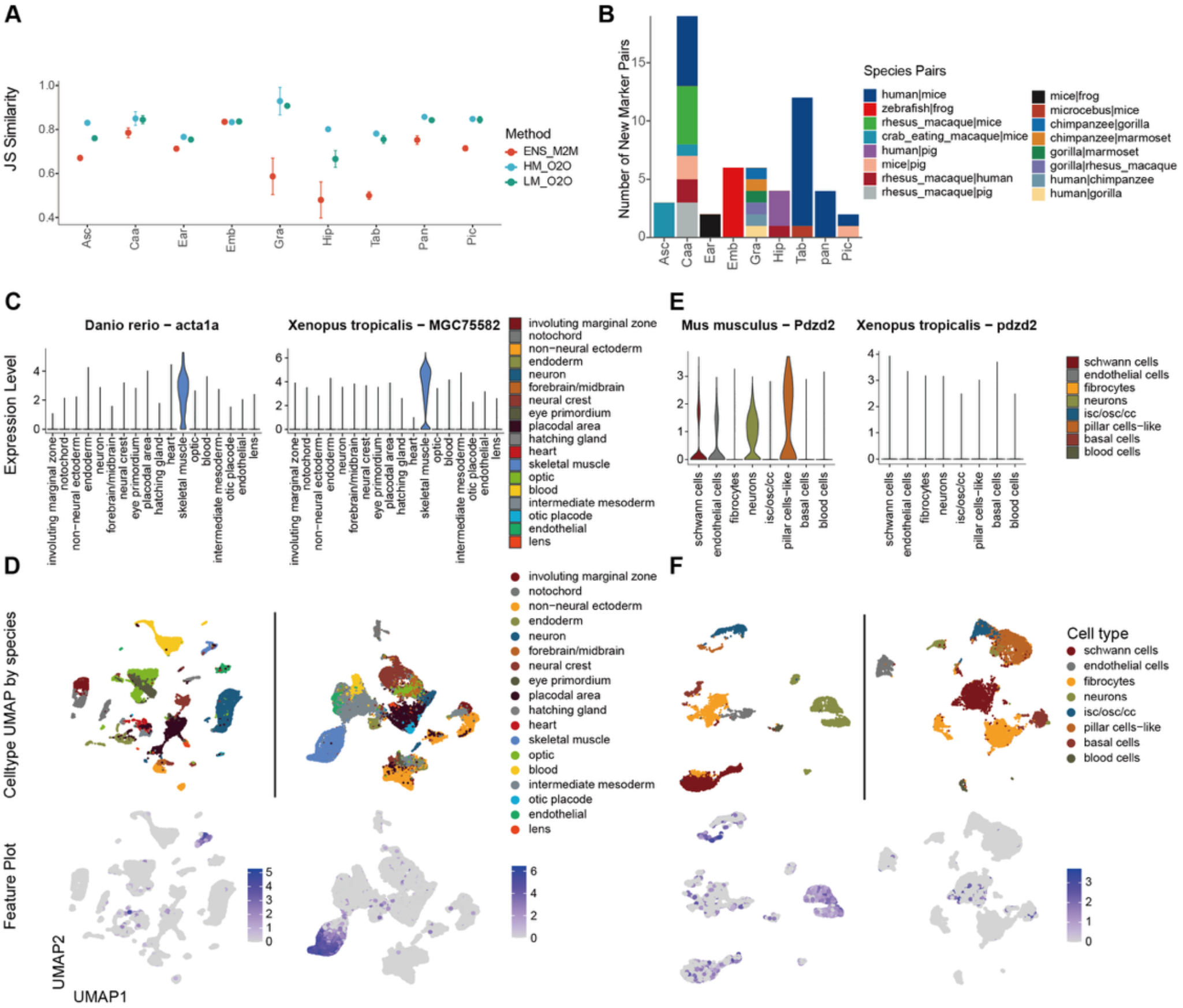
pLLM-based cross-species gene homology mapping uncovers novel marker pairs with high expression concordance. **A**, Jensen-Shannon (JS) similarity of expression profiles for strategy-specific gene homology mappings across nine datasets. Points represent dataset-specific means; error bars indicate mean ± 1.96 × standard error. **B**, Count of novel marker pairs uniquely identified by LM_O2O across species combinations in all benchmarked datasets. **C**, Representative LM_O2O-derived novel marker pair. Violin plots display cross-species expression patterns. **D**, Cellular expression patterns for the marker pair shown in panel C. UMAP visualizations (top) and feature plots (bottom) illustrate their tissue-specific enrichment in *Danio rerio* (left) and *Xenopus tropicalis* (right). **E**, Example of nomenclature-mapped but functionally discordant gene pair annotated in Ensembl. Violin plots reveal divergent expression profiles despite identical gene symbols. **F**, Cellular expression patterns for the gene pair shown in panel E. UMAP visualizations (top) and feature plots (bottom) illustrate their species-specific expression behavior in *Mus musculus* (left) and *Xenopus tropicalis* (right).

We next evaluated the unique contribution of LM_O2O to marker-pair discovery by focusing on LM_O2O-specific gene pairs that were not recovered by sequence-based HM_O2O or Ensembl-based homology annotations. For each species pair across all datasets, we identified LM_O2O-specific gene pairs that were also annotated as cell-type markers (see Methods) (Hao et al. 2024). This analysis identified 58 LM_O2O-specific novel marker-pair occurrences across the nine datasets, corresponding to 54 unique marker pairs, with four pairs detected repeatedly across datasets (Fig. 4B). These pairs provide candidate cases in which embedding-based similarity may capture functional correspondence beyond sequence-based homology. While LM_O2O successfully recapitulated established homologies within closely related gene families (Hansdottir et al. 2004; Bourneuf et al. 2011; Sturm et al. 2014) (Supplemental Fig. 3A), it also proposed previously unannotated correspondences between evolutionarily distant species (Todd et al. 2014) (Supplemental Fig. 3B). For instance, in the Emb dataset, LM_O2O linked Xenopus tropicalis MGC75582(Klein et al. 2002) to Danio rerio acta1a (Ulloa et al. 2015), a pair not recovered by the sequence-based HM_O2O strategy or Ensembl-based homology annotations. Although this correspondence is not apparent from conventional annotation, both genes showed concordant, cell-type-specific expression in skeletal muscle cells, as supported by violin plots and UMAP feature maps (Fig. 4C–D). This example suggests that ESM-derived embeddings can capture functional similarity that is not fully represented by sequence identity or existing homology annotations. To further assess whether these LM_O2O-specific marker pairs represent putative functional correspondences rather than expression-similarity coincidences, we performed protein structure, domain architecture, and external database annotation analyses for the 54 unique marker pairs. Using Foldseek to compare predicted protein structures, 33 of 47 evaluable pairs (70.2%) showed strong, moderate, or local domain-level structural similarity based on qTM-score, tTM-score, and lDDT (Supplemental Fig. S4A, B). We further queried Pfam domains through the InterPro API and found that 44 of 50 evaluable pairs shared at least one Pfam domain, with 16 sharing two or more domains (Supplemental Fig. S4C). To evaluate whether these pairs had been previously annotated in existing orthology resources, we searched NCBI, OrthoDB, and PANTHER. Among the 54 unique pairs, 23 were not recorded in any of the three databases, 11 were supported by one database, 6 were supported by two, and only 14 were recorded in all three databases (Supplemental Fig. S5). Together, the structural and domain-level similarities support the functional plausibility of many LM_O2O-specific marker pairs, whereas the absence of database support for a substantial fraction highlights their potential novelty.

Further joint analysis of expression concordance and marker mapping revealed that shared gene names in Ensembl orthologs do not necessarily reflect functional equivalence. In the Ear dataset, mouse Pdzd2 and frog pdzd2 share an identical gene symbol, yet only the mouse gene exhibits strong, cell-type-specific expression, while its frog counterpart is expressed weekly across cell types (Fig. 4E). Similar gene mappings can also be identified in other datasets (Supplemental Fig. 3C). Taken together, these findings highlight the limitation of sequence homology-based mapping and underscore the complementary value of embedding-based strategies in identifying functionally corresponding genes across species.

## Discussion

Our study presents a systematic evaluation and enhancement of the “gene homology mapping” step in cross-species single-cell transcriptome integration. We demonstrate that one-to-one mapping strategies — whether based on homology (HM_O2O), language model embeddings (LM_O2O), or their fusion (HL_O2O) — consistently outperform the conventional Ensembl many-to-many baseline (ENS_M2M). This conclusion is supported by a comprehensive benchmark spanning 11 species, 9 datasets, and over 3.2 million cells, in which the one-to-one approaches improve key integration metrics (NMI, ARI, cASW, F1 variants, and normalized cLISI), while markedly reducing gene-family–driven noisy micro-clusters. Notably, the language model–based strategy (LM_O2O) not only aligns with classical homology evidence but also complements it by discovering functionally concordant cross-species gene pairs that are previously unannotated.

Gene family expansion and contraction are important evolutionary processes and can encode biologically meaningful species-specific divergence. Therefore, many-to-many homology mappings should not be viewed simply as technical noise. However, in the specific context of cross-species single-cell transcriptome integration, unconstrained many-to-many mappings can mix such biological signals with mapping uncertainty. When one gene in one species is linked to multiple paralogous counterparts in another, expanded gene-family signals may receive disproportionate weight during embedding construction and clustering, producing mapping-associated components or micro-clusters that do not correspond to a single annotated cell type. Our analyses suggest that some ENS_M2M-associated micro-clusters have mixed cell-type composition and lower cross-species expression concordance, consistent with partial mapping-induced bias. At the same time, these observations do not exclude the possibility that a subset of many-to-many-derived signals reflects genuine biology related to paralogous gene families or species-specific divergence. Thus, one-to-one strategies such as HM_O2O, LM_O2O, and HL_O2O should be viewed as approaches for reducing ambiguity in direct cross-species cell-type alignment, rather than as dismissing the biological importance of gene family evolution. In summary, our findings highlight that prioritizing one-to-one mapping and integrating both sequence- and representation-based evidence are pivotal for achieving accurate, interpretable, and generalizable integration of single-cell transcriptome across species. By reducing gene family-driven artifacts while uncovering novel functional gene correspondences, our approach provides a robust and scalable gene-mapping framework to support the construction of cross-species atlases and empower comparative biological discovery.

## Methods

### Implementation of cross-species gene homology mapping strategies

#### Unifying cross-species genes and overall workflow

Robust integration of cross-species single-cell transcriptomics requires unifying genes across species, ensuring that cells are represented in a shared feature space and enabling concatenation of their single-cell matrices.

Based on this principle, we implemented five distinct gene homology mapping strategies, differing in how correspondences are derived and constrained: homology many-to-many (ENS_M2M, derived from Ensemble genes 114), homology one-to-one strict (ENS_O2O, derived from Ensemble genes 114), homology one-to-one (HM_O2O), pLLM–based one-to-one (LM_O2O, derived from ESM2), and a merged scheme combining HM_O2O and LM_O2O (HL_O2O).

Operationally, for each species we begin with raw count matrices and a strategy-specific mapping table, then (i) reindex genes to the mapping set, (ii) assemble a joint matrix over shared genes, and (iii) perform standard single-cell integration steps, including normalization, optional feature selection, and joint embedding. Strategy-specific inputs, mapping rules, and procedures for enforcing one-to-one constraints (e.g., greedy pairing) are provided in subsequent subsections, while metric definitions and evaluation protocols are described elsewhere in Methods.

#### ENS_M2M (many-to-many)

We retrieved pairwise cross-species gene correspondences from Ensembl database (Ensembl Genes release 114). For ENS_M2M, we retained all reported homology pairs between each species pair, thereby preserving the original many-to-many mappings provided by Ensembl.

#### ENS_O2O (one-to-one)

Using the same Ensembl table as ENS_M2M, we restricted correspondences to entries with homology_type = “ortholog_one2one”, yielding a curated one-to-one orthology set for each species pair.

#### HM_O2O (one-to-one)

Starting from the same Ensembl pairwise homology tables as above, HM_O2O enforces a one-to-one mapping by prioritizing Ensembl-derived confidence and sequence identity. For each candidate pair, we extracted three Ensembl attributes: %id. target gene identical to query gene, %id. query gene identical to target gene, and orthology confidence (0=low, 1=high). We defined an identity score (identical_scores) as the mean of the two percent identities. We then applied a global greedy selection over all candidate pairs to resolve many-to-many conflicts: (i) pairs with orthology confidence = 1 are considered before those with 0 (confidence priority); (ii) within the same confidence tier, pairs are sorted by identical_scores in descending order; (iii) iteratively select the highest-priority pair and discard all competing edges incident to either gene, until no conflicts remain. The output is a deterministic one-to-one homology set per species pair.

#### LM_O2O (one-to-one)

For each species pair, we downloaded proteome FASTA files from the Ensembl FTP (release 113) and embedded all protein sequence using the ESM2 protein language model (esm2_t48_15B_UR50D), resulting in a 5120-dimensional representation per sequence. We chose ESM2 because it is a widely benchmarked protein language model that captures structural and functional information and is suitable for large-scale embedding extraction (Schmirler et al. 2024). It provides a practical balance between biological informativeness, ease of use, and computational scalability for the multi-species workflow used here. Protein IDs were mapped to gene symbols via Ensembl annotations. To generate gene-level embeddings from protein isoforms, the workflow supports three aggregation strategies: averaging embeddings across isoforms, dimension-wise max pooling across isoforms, and using only the canonical isoform. In the main analyses, we used averaging as the default strategy because canonical isoform annotations are not uniformly available across all species in our benchmark; among the 11 species included in the nine datasets, only six had canonical isoform annotations available in APPRIS. Averaging across isoforms therefore provides a broadly applicable gene-level representation that is less dependent on the completeness of existing isoform annotations, while the alternative aggregation strategies are available for users with high-quality isoform annotations or species-specific requirements. A cross-species correlation matrix between gene embeddings was computed, and for each gene, the partner with the highest correlation in the other species was selected. The union of selections from both directions defined a double best-hit (DBH) candidate set, which may include one-to-many or asymmetric links. To enforce one-to-one mapping, DBH pairs were ranked by correlation in descending order and a global greedy selection was applied: the highest-ranked pair was iteratively accepted, while all competing pairs involving either gene were discarded, until no conflicts remained. The output is an ESM2-derived one-to-one gene correspondence suitable for integration.

#### HL_O2O (one-to-one)

HL_O2O integrates homology-based and language model–based evidence to generate one-to-one gene correspondences. We first took the union of pairs from HM_O2O and LM_O2O (merging duplicates). For each candidate pair, three attributes were assembled: (i) orthology confidence (from Ensembl; 0/1), (ii) identical_scores (mean of “%id. target gene identical to query gene” and “%id. query gene identical to target gene” from Ensembl), and (iii) correlation (gene-embedding correlation from LM_O2O). If a pair originated from HM_O2O, its correlation was retrieved from the LM_O2O correlation matrix; missing values were imputed with the median correlation of that species pair. Conversely, if a pair originated from LM_O2O, we queried Ensembl for orthology confidence and identical_scores; missing confidence was set to 0, and missing identical_scores were imputed with the median of the candidate set.

After completing attribute collection for all candidate pairs, we applied min–max scaling to normalize the two continuous features (sequence identity and correlation) within each candidate set:

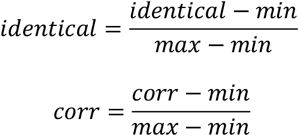

(assigning zero when the denominator was 0). We then computed an unweighted sum to obtain a combined score for each pair:

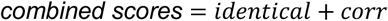

To enforce one-to-one mapping, we applied a global greedy selection procedure with orthology confidence given priority. First, candidate pairs with orthology confidence = 1 were processed before those with 0. Within each confidence tier, pairs were ranked by their combined scores in descending order. The algorithm iteratively accepted the highest-ranked pair and removed all competing edges incident to either gene until no conflicts remained. This procedure produced a deterministic, high-confidence one-to-one mapping that integrates orthology evidence with language model–based similarity.

### Datasets and preprocessing

We analyzed nine datasets, hereafter denoted by abbreviations: Asc (A single cell map of antisense oligonucleotide activity in the brain), Caa (Cell Atlas of Aqueous Humor), Ear (The single cell atlas of inner ear), Emb (Embryo), Gra (Post-mortem cortical tissue of Great apes), Hip (Hippocampus), Tab (Tabula Sapiens, Tabula Microcebus and single cells from different organs of mice), Pan (Pancreas), and Pic (Pancreatic islet cells). All nine datasets were obtained from publicly available cross-species single-cell transcriptomic resources, including the Single Cell Portal, GEO, CZ CELLxGENE, Figshare, and the source publication’s data-availability section. Detailed source links, accession numbers, species composition, and dataset descriptions are provided in Supplemental Table S2.

Dataset selection was based on three practical criteria: availability of public single-cell expression matrices and cell-type annotations for at least two species; coverage of diverse tissues, biological contexts, and evolutionary distances; and inclusion, where possible, of non-model or less commonly represented species. We also avoided over-representing human–mouse comparisons so that the benchmark would not be dominated by the most extensively annotated species pair and could better evaluate the general applicability of LM_O2O across broader evolutionary settings. Datasets were not selected based on the relative performance of LM_O2O or any other mapping strategy.

Quality control (QC) followed the procedures described in the original publications for each dataset. No additional QC was applied, as both dataset-specific and cross-species heterogeneity were substantial, even within individual datasets, making a universal filtering strategy unsuitable for our integration workflow.

Cell-type annotations were adopted from the source studies and standardized across species to maximize one-to-one label correspondence. In cases where species-specific or study-specific nomenclature was used for analogous cell types, we unified them under a consistent naming scheme. Examples include:

#### Caa

schwanncell-nmy → nonmyelinating schwann cell; schwanncell-my → myelinating schwann cell; schwanncell → schwann cell; beamcella, beama → beam a; beamcellb, beamb → beam b; beamx → beam x; beamy → beam y; cribiformjct → jct; bcell → b cell; vascularendo, vascular, vascularendothelium → vascular endothelium; scendo → schlemm’s canal; nkt, nk/t → nk/t cell; mastcell → mast cell; cornealepi, cornealepithelium → corneal epithelium; collectorchnlaqvein → collector channel; pigmentedciliaryepithelium → pigmented ciliary epithelium; nonpigmentedciliaryepithelium → nonpigmented ciliary epithelium; pigmentedepithelium → pigmented epithelium; cornealendothelium → corneal endothelium.

#### Ear

fibrocytes_i, fibrocytes_ii → fibrocytes; marginal stria_i, marginal stria_ii → marginal cells; hair cells_i, hair cells_ii → hair cells; neurons_i, neurons_ii, neurons_iii, type ia neurons, type ib neurons, type ic neurons, type ii neurons, type iii neurons → neurons; pillar cells → pillar cells-like; schwann cells_i, schwann cells_ii → schwann cells.

#### Hip

CA1, SUB → CA1 SUB; CA2, CA3 → CA2-3; Macro, Micro, Myeloid, T → immune; aSMC, vSMC → smooth muscle cell; PC, VLMC → vasculature; COP → OPC; NBs, RGLs, nIPCs → neuronal progenitor.

These harmonizations intentionally reduce label granularity to preserve robust cross-species one-to-one correspondences.

During cross-species integration, to enhance comparability across methods, we retained only cell types that were shared across all species within each dataset. Furthermore, in datasets Asc, Ear, Emb, Gra, Tab, and Pic, cell types with fewer than 100 cells in any species were excluded from all species, ensuring that only cell types with ≥100 cells in every species were kept. This step ensured both reliable sample sizes and consistent cross-species representation.

### Computational requirements and annotation release

ESM2 embedding extraction was performed using esm2_t48_15B_UR50D, which requires approximately 64 GB of GPU memory for large-scale protein embedding generation. To reduce the computational burden for users, we pre-generated gene-level embeddings for the species analyzed in this study and made them available through the GitHub repository described in the Code availability section.

### Biology conservation metrics

To assess the preservation of biological signals in cross-species single-cell transcriptome integration, we employed seven clustering-based evaluation metrics: Normalized Mutual Information (NMI), Adjusted Rand Index (ARI), cell type Average Silhouette Width (cASW), Macro-F1 score, Micro-F1 score, Weighted-F1 score, and Normalized cell-type Local Inverse Simpson’s Index (norm_cLISI). Detailed computational principles of these seven metrics are described below.

*NMI (Normalized Mutual Information)*. Let *U* denote the cluster labels obtained from the integrated embedding and *V* denote the cell-type annotations. With contingency counts *n*_*ij*_ (cells in cluster *i* and cell type *j*) and total sample size *N* = ∑_*ij*_ *n*_*ij*_, we define the empirical frequencies as:

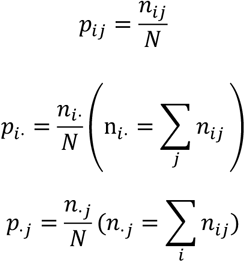

The mutual information and marginal entropies are:

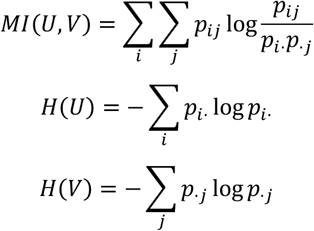

We report the geometrically normalized NMI as:

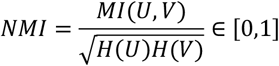

where normalization by the geometric mean of the two marginal entropies symmetrizes and bounds the score. Larger values indicate stronger agreement between clusters and cell types. Unless otherwise specified, logarithms are natural.

#### ARI (Adjusted Rand Index)

The Adjusted Rand Index quantifies the agreement between cluster labels *U* obtained from the integrated embedding and cell-type annotations *V*, corrected for chance. Let *n*_*ij*_ denote the number of cells simultaneously assigned to cluster *i* and cell type *j*, with total sample size *N* = ∑_*ij*_ *n*_*ij*_. Define the marginals as n_*i*_. = ∑_*j*_ *n*_*ij*_ and *n*_·*j*_ = ∑_*i*_ *n*_*ij*_. Using pair counting with the binomial coefficient 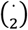, the ARI is given by:

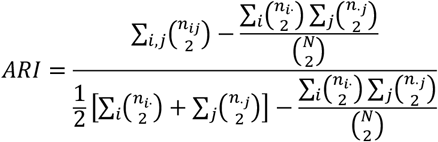

ARI equals 1 for identical partitions, has an expected value of 0 under random labeling, and can be negative when agreement is worse than chance. Its range is [−1,1].

#### cASW (cell-type Average Silhouette Width)

Let *X* ∈ ℝ^*N*×*d*)^ denote the integrated joint embedding, where we used the first 30 dimensions from the Seurat “integrated” reduction obtained via *IntegrateLayers()*, which harmonizes cross-species features prior to downstream analysis. Let *y*_*i*_ denote the cell-type label of cell *i*. Pairwise distances in this space are defined as

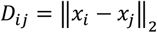

For each cell *i*, define the same-cell-type set

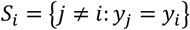

with within-cell-type mean distance

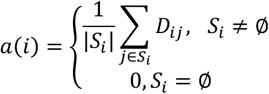

and the nearest other-type mean distance

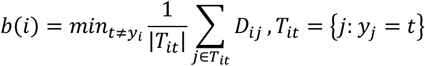

The silhouette value for cell iii is then

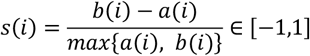

and the cell-type ASW for the dataset is

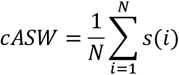

The “integrated” reduction is the low-dimensional space returned by Seurat after cross-species integration via IntegrateLayers(), which harmonizes features across species prior to downstream analysis. and let *y*_*i*_ denote the cell-type label of cell *i*. Pairwise distances are computed in this space (Euclidean by default):*D*_*ij*_ = ‖*x*_*i*_ – *x*_*j*_ ‖_2_. Define the same-cell-type set *S*_*i*_ = {*j* ≠ *i*: *y*_*j*_ = *y*_*i*_} and the within-type mean distance *a*(*i*). *b*(*i*) is the nearest other-type mean distance and *T*_*it*_ denotes a set of all cells whose cell type equals t. For each cell *i*, define

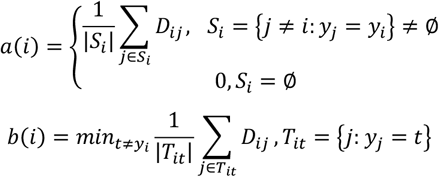

The per-cell silhouette *s*(*i*) is

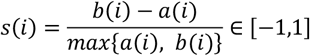

and the cell-type ASW for the dataset is

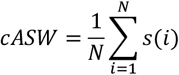

Higher cASW values indicate stronger intra-cell-type cohesion and clearer separation between distinct cell types. Values are reported in the unscaled range [−1,1].

*Macro_F1, Micro_F1 and Weighted_F1*. To evaluate classification performance at both the per-class and overall levels, we computed F1 scores under three aggregation schemes: Macro_F1, Micro_F1, and Weighted_F1. Cluster labels were aligned to cell-type annotations using the Hungarian algorithm applied to the contingency table, and the aligned cluster assignments were treated as predictions ŷof the true labels (*y*). Let the confusion matrix be *C* = [*c*_*uv*_] with rows *u* (true classes) and columns *v* (predicted classes). For each class *t*, we define:

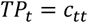

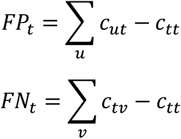

From these, precision, recall, and the per-class F1 score are:

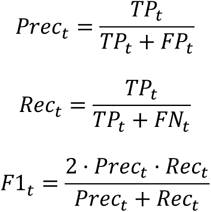

*Macro_F1*. The unweighted mean of per-class F1 scores:

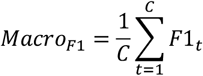

which emphasizes balance across classes, treating rare and abundant types equally.

*Micro_F1*. Computed from globally aggregated counts:

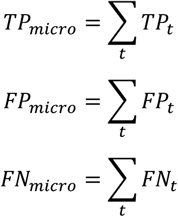

Then

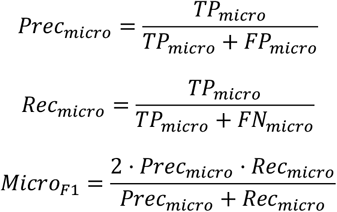

which reflects overall sample-weighted performance, dominated by large classes. Weighted_F1. The support-weighted mean of per-class F1 scores. With class supports *w*, = (∑_*v*_ *cm*_*tv*_)/*N* and ∑_*t*_ *w*_*t*_ = 1, *Weighted*_*F*1_ is calculated as

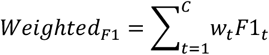

balancing class-specific quality by their prevalence.

Together, these metrics provide complementary perspectives: Macro_F1 highlights per-class balance, Micro_F1 reflects global accuracy, and Weighted_F1 accounts for class size heterogeneity.

*cLISI (cell-type Local Inverse Simpson’s Index)*. cLISI quantifies the preservation of cell-type structure in the integrated embedding by measuring the local purity of cell-type labels within each cell’s neighborhood. Let *K* be the number of distinct cell types and *p*_*it*_ denote the local proportion of type *t* around cell *i*. The per-cell cLISI is defined as the inverse Simpson index:

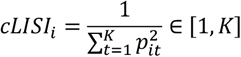

which equals 1 for a perfectly pure neighborhood and increases toward *K* as labels mix. To obtain a conservation-oriented score, we applied geometric normalization to map values into [0,1]:

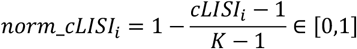

The overall dataset score is the average across all cells:

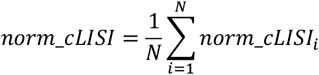

Therefore, a higher *norm*_*cLISI* reflects purer local neighborhoods and improved preservation of cell-type structure after integration.

*iLISI (species Local Inverse Simpson’s Index)*. iLISI quantifies species mixing in the integrated embedding by measuring, for each cell *i*, the local diversity of species label in its neighborhood (as implemented in the R package lisi, function compute_lisi). Let *K*_*sp*_ be the number of species and let *p*_*is*_ denote the local proportion of species *s* around cell *i*. The per-cell iLISI is

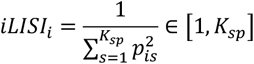

and we report the normalized score as

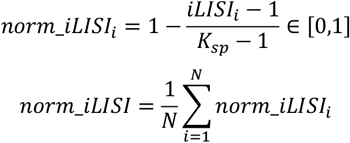

### P-value computation

All p-values in this study were obtained using the Wilcoxon rank-sum test (Mann–Whitney U), as implemented in R (stats::wilcox.test, unpaired).

For the JS similarity comparison between gene homology mapping sources (ENS_M2M vs LM_O2O), we tested the one-sided hypothesis ENS_M2M < LM_O2O using alternative = “less”.

For the correlation comparisons (ENS_M2M vs LM_O2O), we used two-sided tests with alternative = “two.sided” to assess any difference between distributions.

### Bubble heatmap of biology-conservation metrics

Let *D* be the set of datasets (∣ *D* ∣=9) and *K* =

{*NMI, ARI, cASW, Macro*_*F*1_, *Micro*_*F*1_, *Weighted*_*F*1_, *norm*_*cLISI*_} the set of metrics. For method mmm and metric *k* ∈ *K*, we first average the score across datasets:

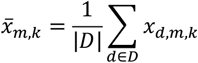

Next, within each metric k we apply min–max normalization across methods (denominator 0 → assign 0.5):

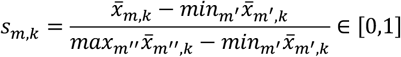

The overall Biology-conservation score (Bioc) for method m is the mean of the seven normalized metrics:

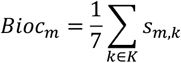

In the bubble heatmap, for non-Bioc cells the bubble radius encodes the normalized score using an area-aware mapping

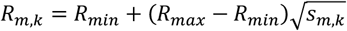

so that bubble area is proportional to *s*_*m,k*_.

Within each metric *k*, methods are ranked in descending order by *s*_*m,k*_; a larger *s*_*m,k*_ implies a better rank (rank = 1 is best). The Bioc column is displayed as a horizontal bar whose length is proportional to *Bio*_*cm*_. The “Mean score” legend refers to the normalized *s*_*m,k*_ scale used for bubble sizing.

### Definition of noisy clusters

After cross-species integration and clustering, we examined the joint embedding and the harmonized cell-type annotations to flag clusters that do not represent a coherent biological cell type. A cluster was labeled noisy if it satisfied all of the following qualitative criteria: (i) it is spatially separated from the canonical cell-type manifolds in the joint embedding (i.e., an outlying micro-cluster); (ii) it contains a mixed composition of cells originating from multiple annotated cell types rather than a single dominant type; and (iii) marker-gene analysis fails to recover a type-specific signature—putative markers show no clear affiliation with any known cell type—such that using this cluster for annotation would likely introduce systematic misassignment. Under this definition, the Pic dataset contained noisy clusters 3, 4, and 5, and the Ear dataset contained noisy clusters 6 and 7 (Fig. 3a–b).

### Source attribution of cross-species gene homology mappings

As introduced before, The HL_O2O catalog is constructed from two upstream sources**—**HM_O2O and LM_O2O. Using explicit record-wise mapping between these two sources, each HL_O2O mapping is assigned to one of three provenance classes: HM_O2O (present only in HM_O2O), LM_O2O (present only in LM_O2O), or HL_COMMON (present in both sources; jointly discovered). We then propagate these pairwise labels to the dataset level by chaining mappings along the dataset’s species order.

### Chaining adjacent species pairs (dataset-level aggregation)

For each dataset, we take the ordered species list and read the mapping for every adjacent species pair. Each pairwise table contains two gene columns and a pair-level source label (HM_O2O, LM_O2O, or HL_COMMON). We successively inner-join neighboring pairs on the shared species to form multi-species chains. For each chained row we compute a combined_property: if all pairwise links share the same label, that label is kept (HM_O2O, LM_O2O, or HL_COMMON); otherwise the row is flagged HL_COMMON.

### ENS_M2M-only set (unique in ENS_M2M mapping strategy)

To identify mappings unique to ENS_M2M, we anti-join the dataset-level HL_O2O table against the dataset-level ENS_M2M table on the species-gene columns, retaining rows absent from HL_O2O. These retained rows constitute the ENS_M2M-only source.

### Filtering to genes present in the dataset

For each species we load the raw feature set (gene names) from HDF5 file of the dataset, standardize to lowercase, and construct a chain key combined_genes by concatenating species-specific gene IDs in the dataset species order. For every source table (HM_O2O, LM_O2O, HL_COMMON, ENS_M2M), we keep only rows whose combined_genes are present for all species (set intersection), then deduplicate. The resulting four source-attributed tables are used in all downstream analyses.

### JS similarity of mapped homology gene expression across species

For each chained mapped homology gene *g*, we quantify cross-species expression similarity at the cell-type level. Given an ordered species list (*s*_1_, …, *s*_*L*_) and a set of cell types *C*, we estimate, separately for every cell type *c* ∈ *C* and species *s*_C_, the probability density of the expression of *g* using a Gaussian KDE on a fixed grid (512 bins) with Silverman’s rule-of-thumb bandwidth *h* = 1.06 σ *n*^−1/5^; densities are computed in PyTorch with a small additive ε = 10^−10^ for numerical stability, yielding probability vectors *p*_*C,l*_.

We then compute the Jensen–Shannon divergence (JSD) between adjacent species and average across the chain. Let the midpoint mixture be

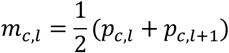

The per–cell-type divergence for gene *g* is

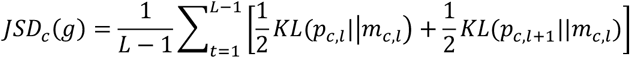

The dataset-level divergence for *g* averages over cell types:

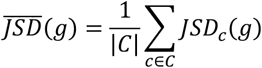

Finally, we report a bounded JS similarity

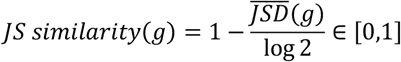

so that larger values indicate more similar cross-species expression distributions for the mapped gene.

### Marker status for mapped homology genes

To assess whether a cross-species mapped homology gene set corresponds to a conserved cell-type marker, we first identified marker genes for each cell type within each original species-specific dataset (prior to integration). Marker genes were defined through differential expression analysis (e.g., using Seurat’s FindMarkers function). For a given dataset and cell type, each species contributes its list of markers. Then, for each mapped homology gene set across species, we checked whether its constituent genes overlap with the marker sets: If all genes in the mapped set are annotated as markers of the same cell type, the pair is classified as a positive marker mapping. If only a subset of the genes is annotated as markers while others are not, the pair is classified as a negative marker mapping. If none of the genes are annotated as markers, the pair is designated as a non-marker mapping. This procedure evaluates whether the derived cross-species correspondences reflect biologically meaningful, conserved marker relationships rather than spurious mappings.

### Protein structural evidence support

To assess structural support for LM_O2O-specific novel marker pairs, we analyzed the 54 unique LM_O2O-specific marker pairs identified by intersecting LM_O2O-specific cross-species gene pairs with cell-type marker genes across all species pairs and datasets. Gene symbols were mapped to UniProt identifiers, and predicted protein structures were obtained from the AlphaFold Protein Structure Database when available. Pairwise structural similarity was evaluated using Foldseek. Seven pairs could not be evaluated because of failed UniProt ID mapping or unavailable AlphaFold structures. For the remaining 47 pairs, we summarized Foldseek structural similarity using qTM-score, tTM-score, mean TM-score, lDDT, fractional identity, and E-value. Structural matches were classified as strong, moderate, local domain-level, or weak based on the combined TM-score and lDDT criteria described in Supplemental Fig. S4.

### Pfam domain evidence support

To evaluate domain-level support for LM_O2O-specific novel marker pairs, genes were mapped to UniProt identifiers and queried against InterPro to retrieve Pfam domain annotations. For each gene pair, we compared the Pfam domains assigned to the two proteins and recorded the number of shared domains. Four pairs could not be evaluated because of unavailable UniProt entries or missing Pfam annotations. Among the remaining 50 pairs, pairs sharing at least one Pfam domain were considered to have domain-level support, and pairs sharing two or more Pfam domains were recorded separately as stronger domain-architecture matches.

### External database support for LM_O2O-specific gene pairs

To assess external support for LM_O2O-specific marker pairs, we searched each candidate cross-species gene pair against three orthology resources: NCBI, OrthoDB, and PANTHER. For each pair, we recorded whether the two genes were supported as homologous or orthologous by each database. Database-support patterns were summarized according to the combination of resources supporting each pair and visualized in Supplemental Fig. S5.

### Cell type annotation transfer accuracy

To evaluate cross-species label transfer, we employed Seurat’s TransferData() function. Anchors were first identified using FindTransferAnchors() on the joint embedding (top 30 dimensions, PCA reduction). Cell type annotations from the reference species were then transferred to the query species via TransferData() (with k.weight = 45).

Accuracy was defined as the proportion of query cells whose predicted labels exactly mapped their ground-truth annotations. To aggregate results across species, we adopted two schemes: If human data were present, human was used as the reference, and each non-human species was treated as a query. The dataset-level accuracy was computed as the cell-count–weighted average of individual query accuracies.

This procedure yields a consistent, size-aware assessment of label transfer across datasets with unbalanced species compositions.

### Three-layer UMAP visualization

To illustrate the process of cross-species cell type annotation transfer, we constructed a three-layer UMAP. All embeddings were generated from the integrated joint space (30 dimensions, PCA reduction, followed by UMAP).

The top layer represents the UMAP of human cells, colored by their annotated cell types. The middle layer shows the UMAP of mouse cells labeled by their predicted annotations, transferred from human using Seurat’s FindTransferAnchors() and TransferData() (with k.weight = 45). Each mouse cell in this layer is connected to its most probable human anchor. The bottom layer displays the UMAP of the same mouse cells but colored by their ground-truth annotations. Each cell in the middle layer is connected vertically to its true annotation in the bottom layer.

To maintain visual clarity, the three layers are vertically offset and only a subsampled set of cells/links is drawn. This design jointly presents the human reference, the transferred predictions, and the true mouse labels within a single figure, providing an intuitive visual assessment of annotation-transfer performance

## Code availability

The code of gene homology mapping strategies and cross_species integration workflow is available at https://github.com/KKzhongyi/pLLM-cross-species-integration. Our scripts from this work are provided as Supplemental Code.

## Acknowledgement

This work was supported by the National Natural Science Foundation of China (32270683 and 32470662); the Beijing Natural Science Foundation (5242006); the Noncommunicable Chronic Diseases-National Science and Technology Major Project (2026ZD0553801); the Fundamental Research Funds for the Central Universities (BMU2021YJ064); We gratefully acknowledge the High-performance Computing Platform of Peking University for conducting the data analyses.

## Author contributions

H.J.W. and Y.J.W. conceived and supervised the project. Z.Y.K. developed and implemented the algorithm, validated the methods, collected and curated the datasets, and wrote the manuscript. Y.C.S. and N.N.W. provided guidance on algorithm development and contributed to manuscript revision. All authors read and approved the final manuscript.

## Notes

### Competing Interest Statement

The authors have declared no competing interest.

### Summary of Updates

Revise and correct the formatting of the manuscript, adjusting the fonts and colors to meet the required guidelines.

