## Supplemental Material for "Protein large language model assisted one-to-one gene homology mapping in cross-species single-cell transcriptome integration"

### Supplemental Figures

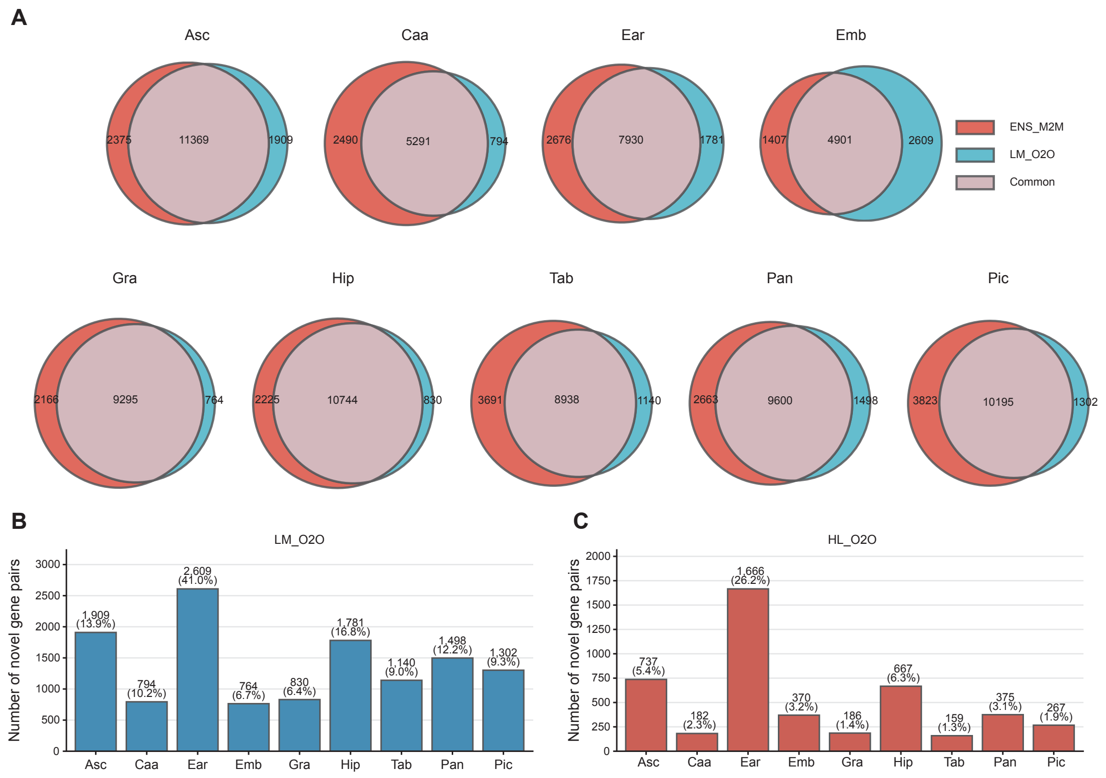


**Supplemental Fig. S1| Comparison of gene-pair overlap and additional one-to-one mappings across homology mapping strategies. A,** Venn diagrams illustrate the overlap of gene correspondences derived from ENS_M2M and LM_O2O across the nine benchmark datasets. **B**, The x-axis shows the nine datasets, and the y-axis shows the difference in the number of gene pairs identified by LM_O2O relative to HM_O2O within each dataset. The number above each bar indicates the absolute difference, and the percentage indicates the proportion of this difference relative to the total number of HM_O2O gene pairs. Because the available expression data for species within each dataset are limited, the observed differences are substantially smaller than the theoretical differences. **C**, Additional gene pairs introduced by HL_O2O across datasets. The x-axis shows the nine benchmark datasets, and the y-axis shows the difference in the number of gene pairs identified by HL_O2O relative to HM_O2O within each dataset. Numbers above the bars indicate absolute differences, and percentages indicate these differences as a fraction of the total number of HM_O2O gene pairs. Because the available expression data for species within each dataset are limited, the observed differences are substantially smaller than the theoretical differences.

**A**

| | Asc | Caa | Ear | Emb | Gra | Hip | Tab | Pan | Pic |
|---|---|---|---|---|---|---|---|---|---|
| Technology | 10x | indrops | 10x | indrops | 10x | 10x | 10x | indrops | 10x |
| Tissue type | Brain | Aqueous humor | Ear | Embryo | Brian | Hippocampus | Cell atlas | Pancreas | Pancreatic islet |
| Shared genes of ENS_M2M | 14622 | 7901 | 11046 | 7229 | 11476 | 13019 | 13074 | 12715 | 14857 |
| Shared genes of ENS_O2O | 13318 | 7630 | 10297 | 5570 | 10548 | 12759 | 11988 | 11921 | 13808 |
| Shared genes of HM_O2O | 13742 | 7781 | 10493 | 6368 | 11459 | 12968 | 12629 | 12263 | 14018 |
| Shared genes of LM_O2O | 13278 | 6085 | 9598 | 7570 | 10059 | 11574 | 10078 | 11098 | 11496 |
| Shared genes of HL_O2O | 14264 | 7929 | 11044 | 7147 | 11770 | 13144 | 12738 | 12554 | 14164 |

**B**

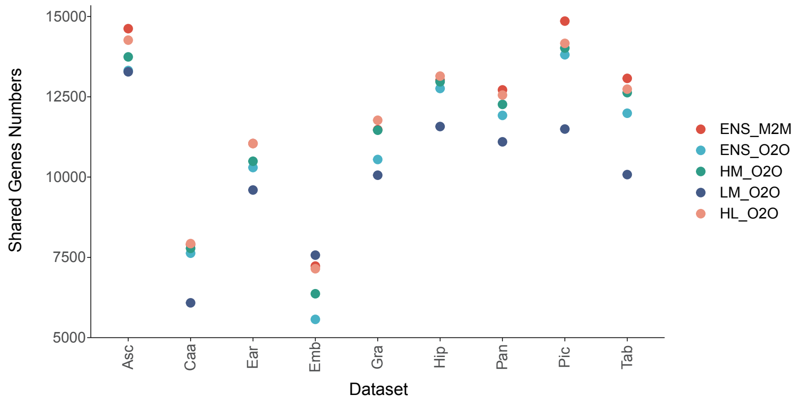


**Supplemental Fig. S2| Dataset characteristics and cross-species gene pair counts across nine datasets. A,** Summary of dataset attributes and gene homology mapping results. Table displays sequencing technology, tissue/organ origin, and the number of cross-species gene pairs identified by each of the five mapping strategies for all nine datasets. **B,** Quantitative comparison of gene pair counts across methods. Scatter plot shows the number of gene pairs per dataset for each strategy.

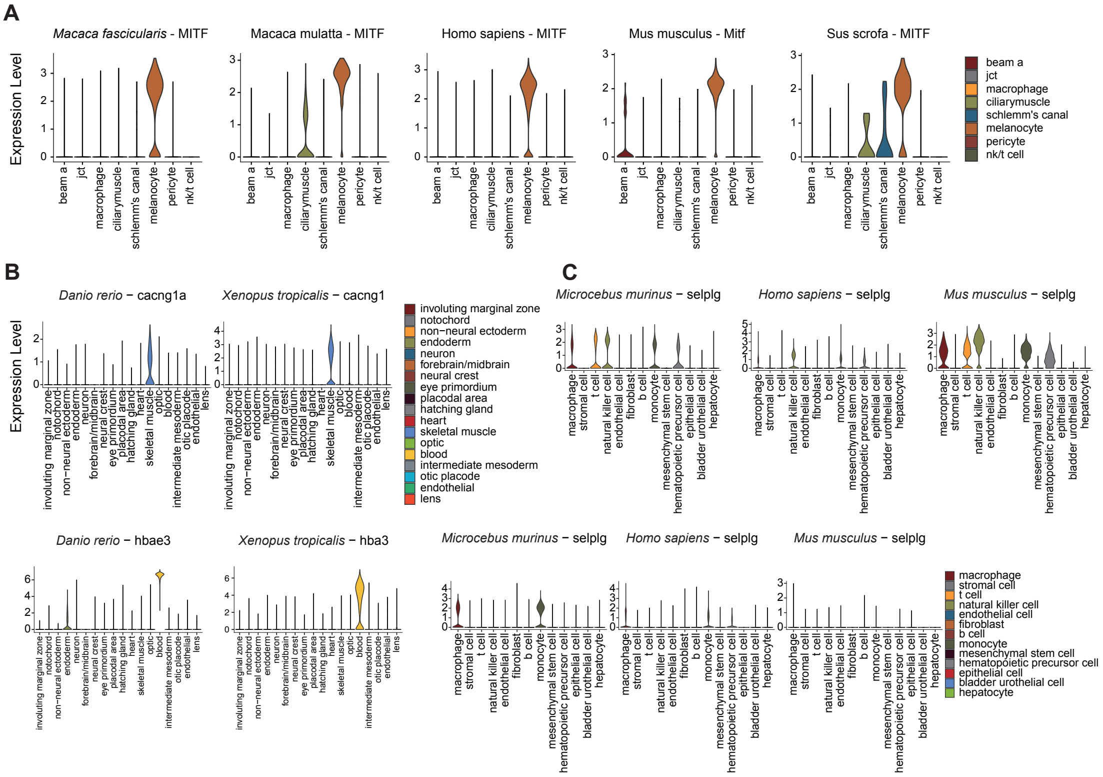


**Supplemental Fig. S3| Examples of marker concordance and discordance in cross-species gene homology mappings. A,** Representative marker pair identified by all methods in Caa dataset. **B,** Representative LM_O2O-derived novel marker pairs. Violin plots display cross-species expression patterns. **C,** Example of nomenclature-mapped but functionally discordant gene pair annotated in Ensembl. Violin plots reveal divergent expression profiles despite identical gene symbols.

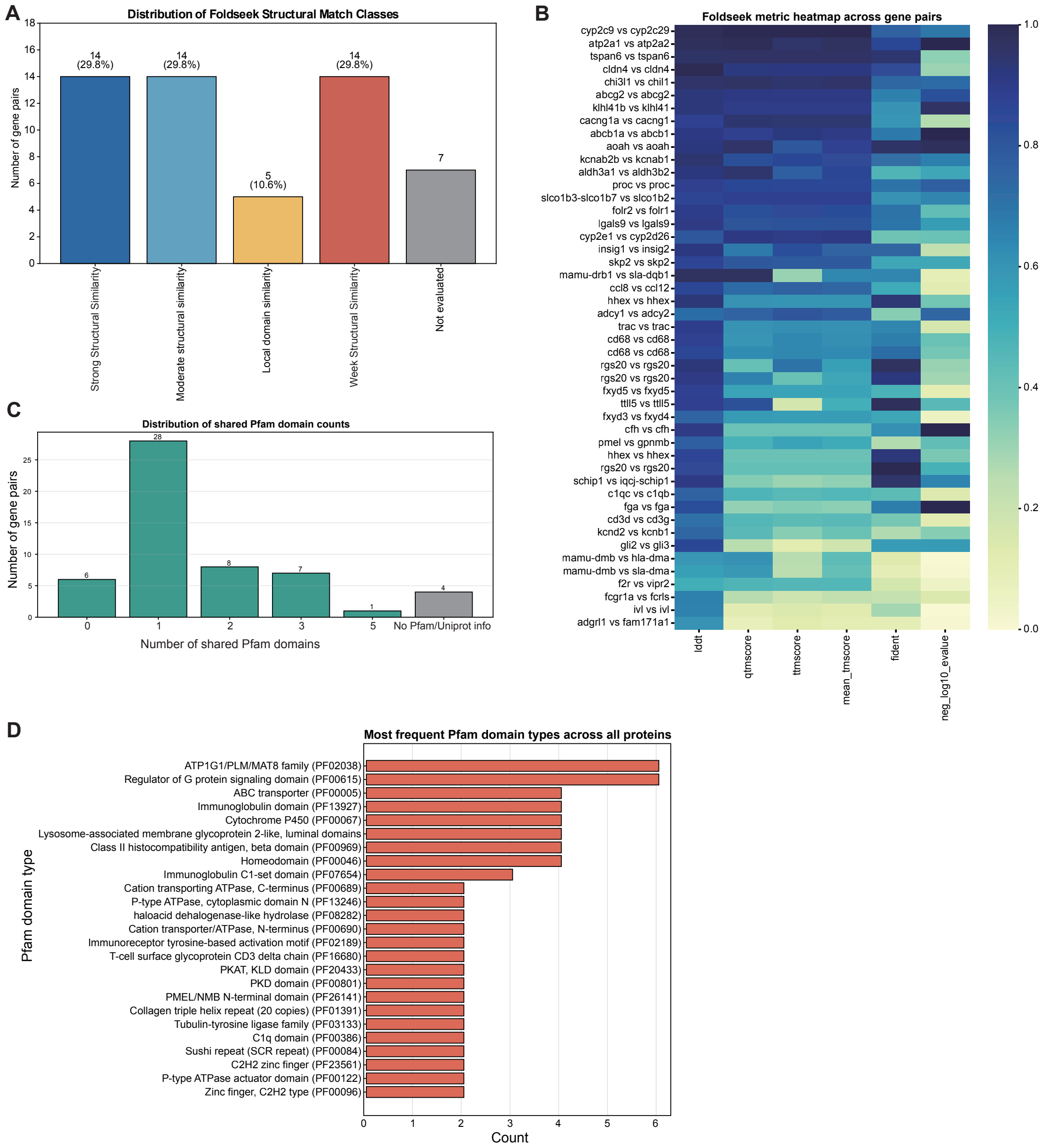


**Supplemental Fig. S4 | Protein structural and Pfam domain evidence supporting putative novel marker gene pairs. A**, Foldseek structural-match classes for the 54 unique putative novel marker-gene pairs. Evaluated pairs were classified as Strong, Local domain, Moderate, or Weak structural similarity. Strong similarity required high TM-scores in both query- and target-normalized directions [min(qtmscore, ttmscore) ⩾ 0.80] and lDDT ⩾ 0.80. Local domain similarity was assigned when similarity was mainly localized to a domain and met any of the following criteria: max(qtmscore, ttmscore) ⩾ 0.60, min(qtmscore, ttmscore) < 0.50, and lDDT ⩾ 0.65; mean_tmscore ⩾ 0.50, |qtmscore − ttmscore| ⩾ 0.30, and lDDT ⩾ 0.70; or lDDT ⩾ 0.75 and max(qtmscore, ttmscore) ⩾ 0.45. Moderate similarity was assigned to pairs that failed the local-domain criteria but had mean_tmscore ⩾ 0.50 and lDDT ⩾ 0.70; all remaining pairs were classified as Weak. **B**, Foldseek-based comparison of predicted protein structures for the same 54 pairs. Rows indicate gene pairs and columns indicate Foldseek metrics, including lDDT, qTM-score, tTM-score, mean TM-score, fractional identity, and neg_log10_evalue. lDDT measures local structural consistency; qTM-score and tTM-score are TM-scores normalized by query and target protein length, respectively; mean_tmscore is their average; fident is the fraction of identical residues in the aligned region; and neg_log10_evalue is −log10(E-value), indicating Foldseek hit significance. For heatmap visualization only, neg_log10_evalue was min–max normalized from its original range of 2.448184–94.677784; all other Foldseek metrics are shown as original values. **C**, Distribution of shared Pfam domains across the 54 putative novel marker-gene pairs. **D**, Most frequent Pfam domain types detected across proteins in the putative novel marker-pair set, ranked by occurrence count.

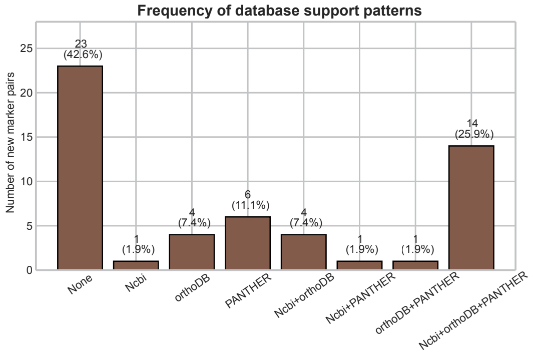


**Supplemental Fig. S5 | Database support patterns for 54 functionally concordant cross-species gene pairs.** Bar plot summarizing the database support patterns for 54 functionally concordant cross-species gene pairs identified by LM_O2O, based on searches in three orthology resources: NCBI, OrthoDB, and PANTHER. Each bar shows the number of gene pairs supported by a specific combination of databases, with the corresponding percentage indicated above the bar.

Mean clustering metrics across nine datasets for five cross-species matching methods

| Method | NMI | ARI | norm_cLISI | Macro-F1 | Micro-F1 | Weighted-F1 | cell_type ASW |
|---|---|---|---|---|---|---|---|
| ENS_M2M | 0.681481 | 0.586384 | 0.984838 | 0.648577 | 0.726251 | 0.720203 | 0.271135 |
| ENS_O2O | 0.773793 | 0.703580 | 0.984149 | 0.745534 | 0.795105 | 0.802900 | 0.277389 |
| HM_O2O | 0.791681 | 0.730563 | 0.984724 | 0.762503 | **0.810883** | **0.817959** | 0.277691 |
| LM_O2O | 0.783729 | 0.719947 | 0.984385 | 0.761391 | 0.802260 | 0.810110 | 0.271576 |
| HL_O2O | **0.797985** | **0.734940** | **0.985763** | **0.764334** | 0.809052 | 0.810631 | **0.281400** |

**Supplemental Table S1 | Summary table showing the mean performance of five cross-species gene-mapping methods across nine benchmark datasets and seven clustering metrics.** Bold values indicate the best-performing method for each metric.

**Supplemental Table S2 | Public data sources used in this study.** Summary of the nine publicly available cross-species single-cell transcriptomic datasets used for benchmarking. The table includes dataset abbreviation, tissue or biological context, species composition, data source, accession or collection information, and source URL for each dataset (see the Excel table).
