## Supplemental_Figures for "Protein large language model assisted one-to-one gene homology mapping in cross-species single-cell transcriptome integration": Supplemental_Fig_S2.pdf

**A**

| | Asc | Caa | Ear | Emb | Gra | Hip | Tab | Pan | Pic |
|---|---|---|---|---|---|---|---|---|---|
| Technology | 10x | indrops | 10x | indrops | 10x | 10x | 10x | indrops | 10x |
| Tissue type | Brain | Aqueous humor | Ear | Embryo | Brian | Hippocampus | Cell atlas | Pancreas | Pancreatic islet |
| Shared genes of ENS_M2M | 14622 | 7901 | 11046 | 7229 | 11476 | 13019 | 13074 | 12715 | 14857 |
| Shared genes of ENS_O2O | 13318 | 7630 | 10297 | 5570 | 10548 | 12759 | 11988 | 11921 | 13808 |
| Shared genes of HM_O2O | 13742 | 7781 | 10493 | 6368 | 11459 | 12968 | 12629 | 12263 | 14018 |
| Shared genes of LM_O2O | 13278 | 6085 | 9598 | 7570 | 10059 | 11574 | 10078 | 11098 | 11496 |
| Shared genes of HL_O2O | 14264 | 7929 | 11044 | 7147 | 11770 | 13144 | 12738 | 12554 | 14164 |

**B**

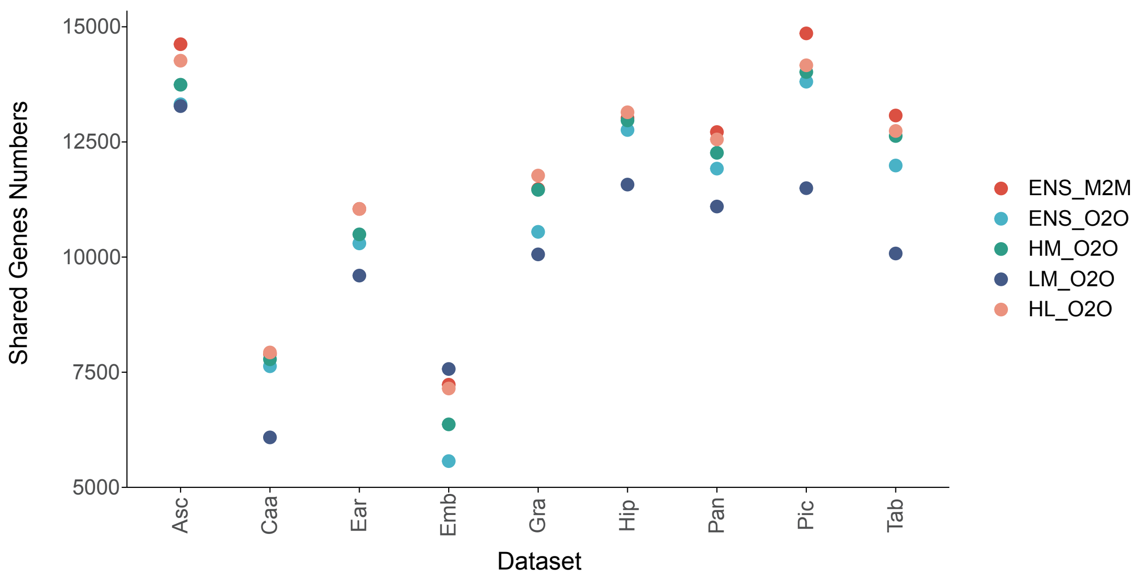
