## Supplementary figures and images for "Protein large language model assisted one-to-one gene homology mapping in cross-species single-cell transcriptome integration"

### Supplemental_Fig_S1.pdf

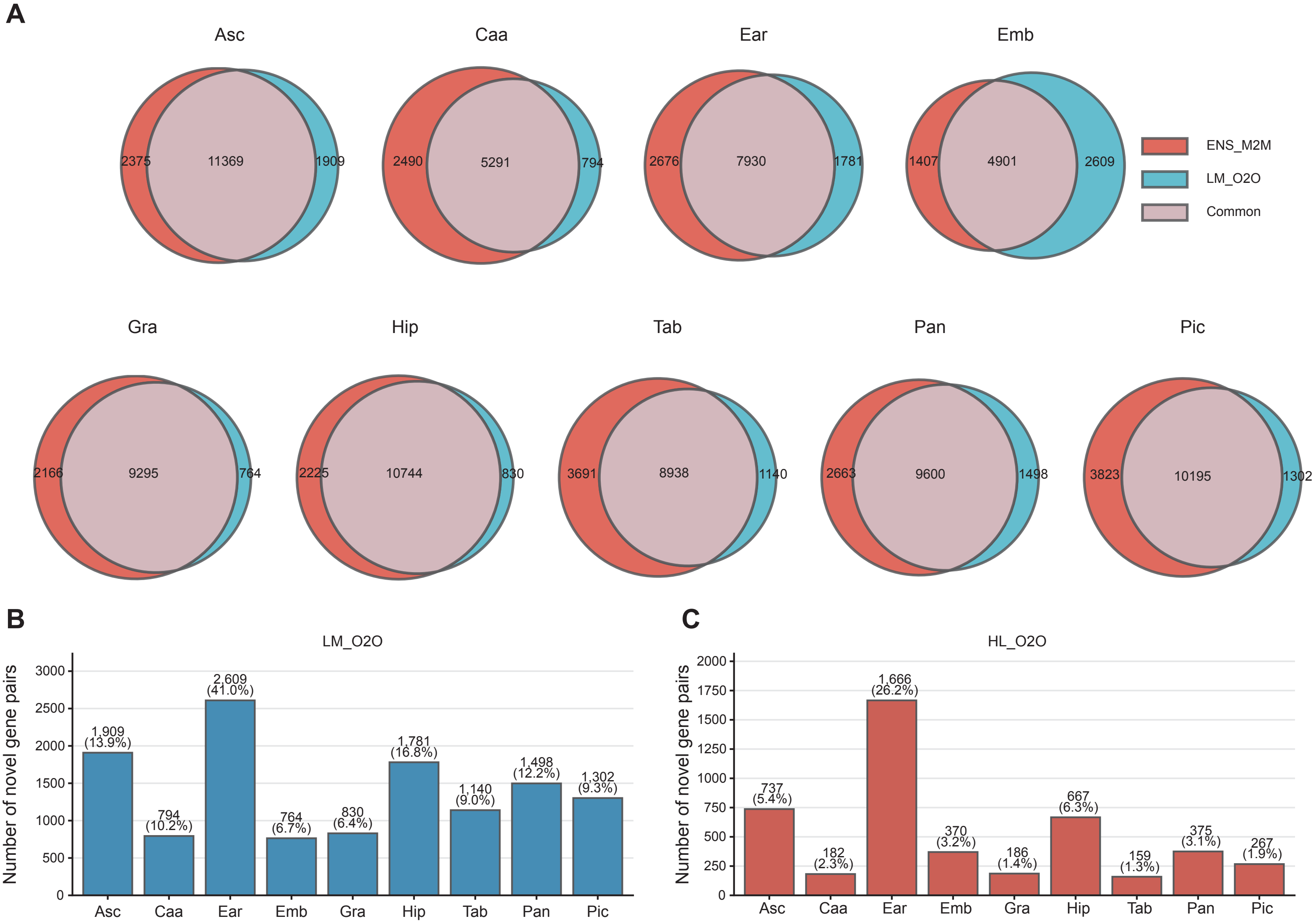

### Supplemental_Fig_S3.pdf

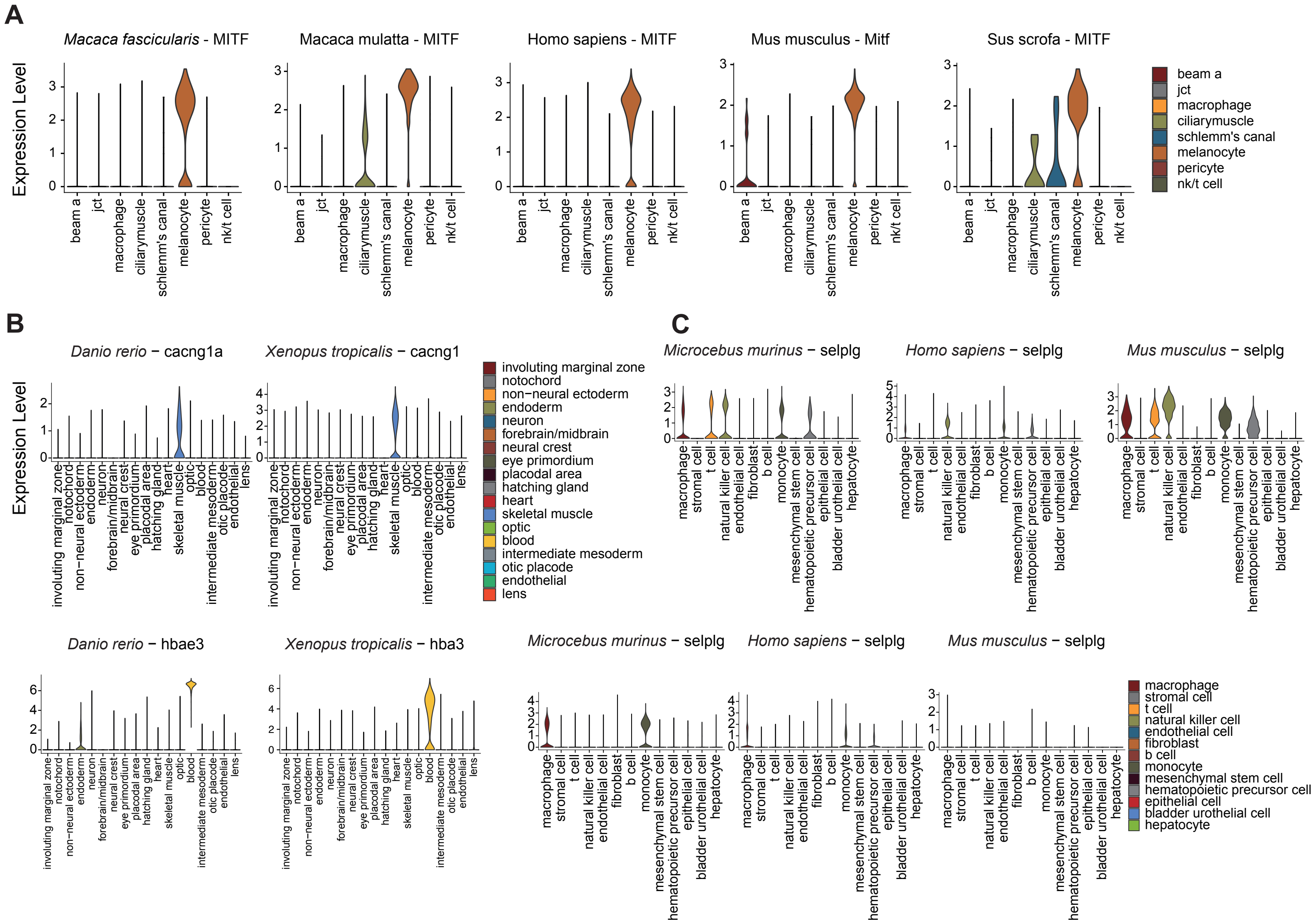

### Supplemental_Fig_S4.pdf

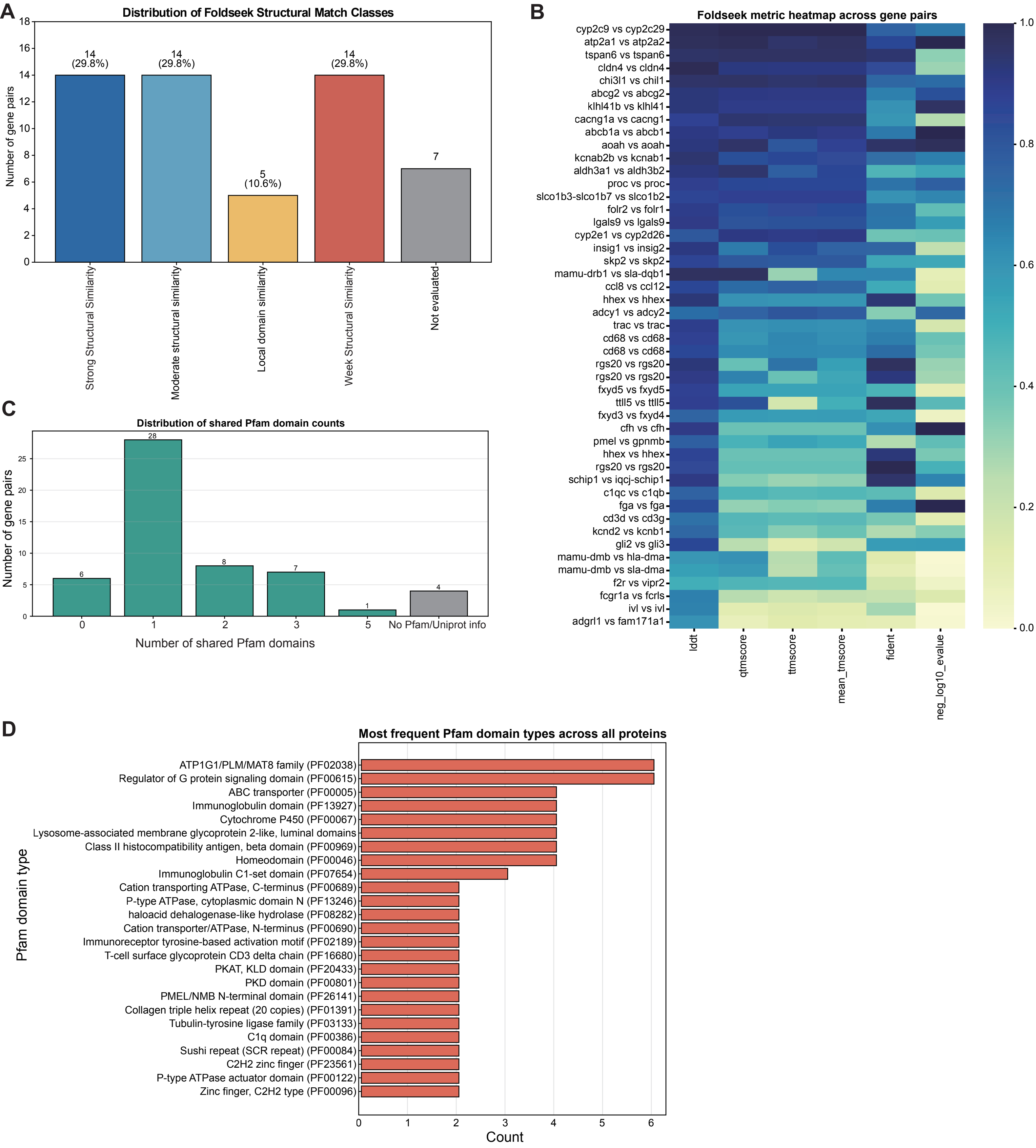

### Supplemental_Fig_S5.pdf

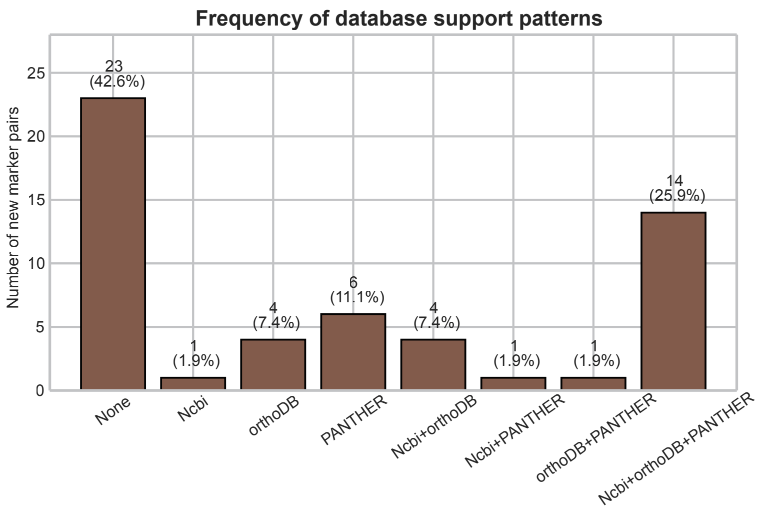
